# Mechanical stimulation of endocardial Purkinje fibres can trigger ventricular arrhythmias

**DOI:** 10.1101/2020.08.10.234195

**Authors:** Ed White, Richard Walton, Sarbjot Kaur, Amelia Power, Michel Haïssaguerre, Olivier Bernus, Marie-Louise Ward

**Affiliations:** School of Biomedical Sciences, University of Leeds, Leeds, UK; Department of Physiology, University of Auckland, Auckland, New Zealand; IHU Liryc, Electrophysiology and Heart Modeling Institute, fondation Bordeaux Université, F-33600 Pessac-Bordeaux, France; Univ. Bordeaux, Centre de recherche Cardio-Thoracique de Bordeaux, U1045, F-33000, Bordeaux, France; INSERM, Centre de recherche Cardio-Thoracique de Bordeaux, U1045, F-33000 Bordeaux, France; Bordeaux University Hospital (CHU), Electrophysiology and Ablation Unit,F-33600 Pessac, France

**Author notes:** University of Otago, Dunedin, New Zealand (Amelia Power).

## Abstract

Acute ventricular dilation can evoke mechanically-induced arrhythmias, our study investigated whether Purkinje fibres (PFs) may play a role. Changes in left ventricular (LV) pressure and pseudo-ECGs were measured in isolated, Langendorff-perfused, male Wistar rat hearts in sinus rhythm. The LV endocardial surface was irrigated with experimental agents, via an indwelling catheter. Mechanically-induced arrhythmias were triggered by LV lumen inflation (100μl in 2s) via an indwelling balloon. Arrhythmias occurred as the LV volume was increased and spontaneously ceased within 20s of the onset of LV inflation. Arrhythmias were indexed as an increase in the standard deviation of all R-R intervals (SDRR), the number of ectopic activations and the period of these activations. Following 10s LV endocardial irrigation with Lugol’s solution (IK/I_2_) to chemically ablate surface PFs or with 0 Na^+^ Tyrode, there was a statistically significant attenuation of mechanically-induced arrhythmias. Lugol’s reduced the mechanically-induced increase in SDRR (Tyrode pre-stretch 3.5±1.7ms to 113.8±15.1ms during stretch vs Lugol pre-stretch 3.3±0.5ms to 39.9±14.5ms during stretch n=8, P < 0.05). There was also a reduction in the number (21.2±2.0 to 1.5±0.7, P<0.001) and period (5.9±0.71s to 1.7±0.85s, P< 0.01) of ectopic activations. The experiment was repeated using LV lumen irrigation with either 1µM GsMTx4, a peptide that blocks stretch-activated channels or 50µM 9-Phenanthrol (9-Phen), a blocker of TRPM4 channels. GsMTx4 did not attenuate mechanically-activated arrhythmias while 9-Phen had a partial effect. 9-Phen statistically reduced the number and period of ectopic activations but did not attenuate the mechanically-induced increase in SDRR (n=6-11 for each intervention). In further studies, *in situ* focal mechanical stimulation of individual PFs, caused ectopic activations, in each of 4 sheep LV preparations (0.2±0.1 ectopics in 10s pre-mechanical stimulation vs 2.1±0.2 ectopics in 10s during mechanical stimulation, P<0.001). We interpret our observations in rat and sheep hearts as evidence for a role of PFs in the generation of some mechanically-induced arrhythmia.

## Introduction

### Mechanical stimulation in the heart and mechanically-induced arrhythmias

The mechanical environment of the heart is important in both normal and pathological function. Changes in cardiac preload and afterload and associated alteration of chamber volume result in the modulation of sinus rhythm via stretch of the sino-atrial node (the Bainbridge effect), contractility via stretch of the atrial and ventricular myocardium (the Frank-Starling effect) and action potential configuration (Quinn & Kohl, 2016).

The inter-dependence of mechanical and electrical activity is demonstrated by the processes of excitation-contraction coupling (Bers, 2002) and mechano-electric feedback/coupling (Orini *et al*., 2017; Quinn & Kohl, 2020). A manifestation of these interactions are mechanically-induced arrhythmias which occur in several clinical settings such as atrial fibrillation, *commotio cordis*, hypertrophy and heart failure e.g. (Kohl *et al*., 2011;Orini *et al*., 2017;Yamazaki *et al*., 2012; Quinn, 2014).

There is strong evidence that the myocardium can be the source of mechanically-induced arrhythmias e.g. (Quinn *et al*., 2017) showed that in isolated rabbit hearts, focal mechanical stimulation of the epicardial surface could trigger arrhythmias arising from the site of mechanical stimulation. Mechanically-induced arrhythmias are typically explained in terms of the activation of stretch-activated, non-specific cationic ion channels within the myocardium, though the identity of these in the ventricular myocardium is uncertain (White, 2006;Syeda *et al*., 2015; Quinn & Kohl, 2020).

A common experimental technique used to stretch *in situ* and intact, isolated hearts, is chamber lumen expansion. This is achieved, either by using an indwelling fluid-filled balloon (Franz *et al*., 1992), by increasing filling pressure or by briefly occluding chamber outflow. These techniques provoke mechanically-induced arrhythmias in many species including rat (Kim *et al*., 2012); rabbit (Franz *et al*., 1992); sheep (Chen *et al*., 2004) and human (Levine *et al*., 1988).

Purkinje fibres (PFs) line the endocardial surface and will be stretched by lumen inflation (Canale *et al*., 1983). Thus, mechanical activation of PFs is a plausible aspect of mechanically-induced arrhythmias (Reynolds et al, 1975; Ferrier, 1976; Thakur et al, 1996), additional to the activation of the myocardium. This interpretation has not been extensively tested experimentally, even though earlier work on excised PFs presented evidence that they are sensitive to mechanical manipulation (Dudel & Trautwein, 1954; Kaufmann & Theophile, 1967;Deck, 1964;Dominguez & Fozzard, 1979).

### PFs and arrhythmias

PF structure, Ca^2+^-handling and electrical properties are distinct from the ventricular myocardium, see (Boyden *et al*., 2016;Dun & Boyden, 2008;Boyden *et al*., 2010;Haissaguerre *et al*., 2016). PFs are implicated in the induction and maintenance of arrhythmias, including monomorphic and polymorphic ventricular tachycardia and ventricular fibrillation (Haissaguerre *et al*., 2002;Haissaguerre *et al*., 2016;Samo *et al*., 2014). These arrhythmias are prevalent after myocardial infarction, possibly due to slowed conduction or depolarisation of ischemic tissue (Haissaguerre *et al*., 2016). Triggered, ectopic foci with early after depolarisation or delayed after depolarisation forms may be created by dysfunctional Ca^2+^ handling (Haissaguerre *et al*., 2016;Boyden *et al*., 2016) and/or dysfunctional transient outward K^+^ current (Boyden *et al*., 2016), with ventricular tachycardia maintained by re-entry within the PFs (Haissaguerre *et al*., 2016). PF ablation can be an effective treatment for these arrhythmias (Haissaguerre *et al*., 2018;Nogami, 2011b;Nogami, 2011a). Despite the importance of PFs in arrhythmia initiation and maintenance, mechanisms of action are not fully understood (Haissaguerre *et al*., 2016) thus a better knoweledge of the role of mechanical stimulation in PF arrhythmias is important.

Our study tested the hypothesis that PFs can play a role in the generation of mechanically-induced arrhythmias. We used isolated rat hearts to study the effect of stretch on intact hearts. As these studies did not allow us to verify directly the stimulation of PFs, we also performed studies in a larger species, sheep, that allowed us to identify and target individual PFs for mechanical stimulation and to record activation via optical mapping.

## Methods

Wistar rats (N = 19, 250-280g) were killed in accordance with local and national ethical regulations and approvals, University of Auckland AEC 001232, UK Home Office PPL No 70/8399. Hearts were rapidly removed and Langendorff perfused with a HEPES buffered physiological saline at 38°C that contained (in mM): NaCl 142; KCl 6; MgSO_4_ 1.2; Na_2_HPO_4_ 1.2; HEPES 10, Glucose 10; CaCl_2_ 1.8. (pH 7.4). All chemicals were purchased from Sigma except GsMTx4 which was purchased from Peptide International Inc, USA.

A deflated cellophane balloon (connected to a pressure transducer and fluid filled syringe) and another tube (outer diameter, 0.9mm) were placed in the left ventricular (LV) lumen via the mitral valve and an opening created by the removal of part of the left atria and secured in place by tying to the aortic cannula. The opening of the mitral valve and removal of left atrial tissue provided a route for LV drainage. Thus, the myocardium was perfused via the aorta while the tube in the LV lumen was used to irrigate the LV endocardial surface. Needle electrodes were used to record pseudo-ECGs (negative lead in the right atria, positive lead in the ventricular apex avoiding the ventricular lumens).

The deflated balloon was inflated until a stable pressure pulse was detected, it was assumed this indicated the balloon was in contact with the LV endocardial surface. The LV was stretched by fluid inflation of the balloon with an additional 100µl over approximately 2s. Balloon inflation was maintained for at least 30s (typically 40s). The balloon was then deflated by withdrawing 100µl. Thus, in a given heart the LV was stretched to and from constant volumes. This stretching protocol was applied before and after washing of the endocardium with solutions introduced into the LV lumen via the indwelling tube. The inflation speed and volume of the balloon was consistent with those we have previously used to trigger mechanically-induced arrhythmias in hearts from similar sized animals (Benoist *et al*., 2014). These were deemed appropriate for this study as they reproducibly provoked mechanically-induced arrhythmias, allowing us to investigate the phenomenon.

Either 1ml of Tyrode or (i) 0.1ml Lugol’s solution (120mM KI, 38.6 mM I_2_) over 10s followed by 0.9ml Tyrode (ii) 3ml of 1µM GsMTx4 over 5 min (iii) 3ml of 50µM 9-Phenanthrol (9-Phen) over 5min (iv) 0 Na^+^ Tyrode, irrigated the endocardial lumen via the indwelling tube. In each heart, 2 stretches, approximately 60s apart, were performed immediately following Tyrode irrigation and 2 immediately following the intervention (see supplementary Figure S1).

Pseudo-ECG records were used to identify sinus and ectopic activations. Intrinsic heart rate was calculated from R-R intervals of the pseudo-ECG in sinus rhythm. The standard deviation of R-R intervals (SDRR), including sinus and ectopic activations, was used as one measure of stretch-disruption to sinus rhythm. Additionally, the number of non-sinus activations caused by stretch (ectopic activations) was measured. The period of ectopic episodes was also measured, as the time from the first ectopic to the time of the first sinus beat following the last ectopic. Ectopic activations were identified by their timing and waveform, relative to sinus rhythm activations. Data was analysed from 20s of continuous records immediately before stretch (Pre) and 20s from the beginning of stretch (Stretch), this 20s period captured all ectopic activity (see Results).

Changes in pressure (rather than absolute pressures) were used to indicate stretch via increases in LV developed pressure (LVDP) and diastolic pressure (δDiaP) and on the impact of pharmacological interventions on LVDP prior to stretch. LVDP was the difference between diastolic and systolic pressures, δDiaP was the difference in diastolic pressures at the end of each pre-stretch and during-stretch 20s sampling period. Data was acquired at 1 kHz and analysed with Labchart (ADInstruments, NZ). All data were collected in sinus rhythm at 38°C.

Brief exposure to Lugol’s solution has previously been reported to destroy surface and free-running PFs but leave the underlying myocardium largely intact e.g. (Chen *et al*., 1993). This intervention has been used to study the role of PFs in arrhythmias e.g. (Dosdall *et al*., 2008;Myles *et al*., 2012). GsMTx4, is a peptide known to block non-specific cationic stretch-activated ion channels (Bode *et al*., 2001;Bowman *et al*., 2007), 9-Phen is a blocker of TRPM4 channels, which modulate PF electrical activity (Hof *et al*., 2016;Guinamard *et al*., 2015). Dosage was based on previously reported effects. 0 Na^+^ Tyrode was used to inhibit phase 0 activation, solution tonicity was maintained by replacing NaCl with 284 mM sucrose.

Sheep studies were carried out as previously described (Martinez *et al*., 2018) and in accordance with the recommendations of the Directive 2010/63/EU of the European Parliament on the protection of animals used for scientific purposes and the local ethical committee of Bordeaux CEEA50. Hearts were obtained from female adult sheep (N = 4; 42-55kg). Sheep were pre-medicated with ketamine (20mg/kg) and acepromazine (Calmivet, 1mL/50 kg). Anesthesia was induced with intravenous injection of sodium pentobarbital (10 mg/kg) and maintained under isofluorane, 2%, in 100% O_2_. Sheep were euthanized by sodium pentobarbital (40 mL, from 50 mg/mL stock) and the heart rapidly excised, cannulated by the aorta and rinsed with cold cardioplegic solution, containing (mM): NaCl, 110; CaCl_2_, 1.2; KCl, 16; MgCl_2_, 16; NaHCO_3_, 10 and glucose, 9.01 at 4°C. The LV wall was dissected and cannulated by the left anterior descending and circumflex arteries. LVs were mounted on a frame and were submersed and perfused (20mL/min) with a warm (37°C) saline solution containing (mM): NaCl, 130; NaHCO_3_, 24; NaH2PO_4_, 1.2; MgCl_2_, 1; glucose, 5.6; KCl, 4; CaCl_2_, 1.8; gassed with 95% O_2_/5% CO_2_ at 37°C (pH 7.4); and supplemented with blebbistatin (10µM), to inhibit contraction.

Ventricles were stained with the potentiometric dye di-4-ANEPPS (10µM, delivered via the cannulated arteries and recirculation) which was excited by illumination of the endocardial surface using monochromatic LEDs at 530 nm (Cairn Research Ltd, Kent, UK). A Micam Ultima CMOS camera (SciMedia USA Ltd, CA, USA) with a 100×100 pixels resolution was used to obtain images at a spatial resolution of 850µm/pixel and a frame rate of 1kHz. Optical emission signals were selected using a broad-band 650 ± 20nm filter.

PF were chosen that did not span the LV lumen, to avoid PFs that might have been over extended by the opening of the LV. PFs were not adhered to the endocardial surface other than by their terminal insertion points. LV preparations were electrically paced via bipolar Tungsten microelectrodes with a resistance of 2MOhms. The electrodes were insulated by a 3μm kapton sheath (electrode OD 0.356mm) exposing a tip of length 5mm and tip diameter of 1-2μm (World Precision instruments, ref TM33B20KT). They were placed either in the endocardial surface, or in a chosen PF (Martinez *et al*., 2018). Preparations were stimulated at 0.5Hz with 5ms pulses which were increased in amplitude until regular capture was achieved. For PF stimulation the two electrode tips were inserted into the PF collagen sheath along its longitudinal axis with an inter-electrode spacing of approx. 1mm, ensuring no contact with the surrounding myocardium. PF electrode sites were away from the fibre root with the myocardium.

PFs were mechanically stimulated by contacting a single PF with a 2mm diameter wooden probe (6-10 contacts over a 10s period). The individual contacts were applied manually, were as brief as physically possible and of a pressure that did not damage the PFs (as evidenced by the ability of the chosen PF to respond to electrical stimulation following the mechanical stimulation). When this stimulus was replicated by contacting an inflated balloon connected to a pressure transducer, contact produced a pressure transient of 32.7±2.5 mm.Hg which took 86.1±6.1ms to peak and had a total duration of 262.6±2.6 ms (N = 8 trails of 10 stimuli per trail). The start and end of the period of mechanical stimulation in the sheep LV was recorded but timing, relative to the electrical stimulus was unknown.

Following mechanical stimulation of a PF, the stimulating electrodes were transferred from the endocardium to the PF in order to compare activation patterns and the viability of the PF, post-mechanical stimulation. Ectopic activations were identified as activations not associated with the external trigger stimulus. Pseudo-ECGs were recorded and analysed with Labchart. Optical signals were analysed with BV Analysis (Brainvision) to identify the location and timing of earliest activation.

### Statistical analysis

Data were analysed with SigmaStat 3.5. Effects of interventions were tested by paired Student’s t-test with the exception of the effect of treatments and stretch on SDRR. These were tested by 2-way repeated measures analysis of variance (2RMANOVA) with treatment and stretch as factors for each heart. Corrected pairwise post-hoc comparisons were made by Holm-Sidak tests following a statistically significant ANOVA. The term ‘significantly’ in the text refers to a statistically significant effect (P <0.05).

## Results

Left ventricular stretch by balloon inflation generated ectopic activations that caused a disruption of the rhythmic pattern of pseudo-ECG R-waves and associated systolic pressure transients (Fig.1). Ectopic excitations manifested several morphologies with both +ve and – ve vectors. Fig. 1 demonstrates isolated ectopics, doublets and brief tachycardia, relative to sinus rhythm. Episodes of ectopic activity began as LV balloon volume was increased and were most frequent in the ensuing 5s. The mean period of ectopic activity is given in Table 1. Sinus rhythm spontaneously returned in all preparations within 20s of the start of stretch.

**Table 1.**
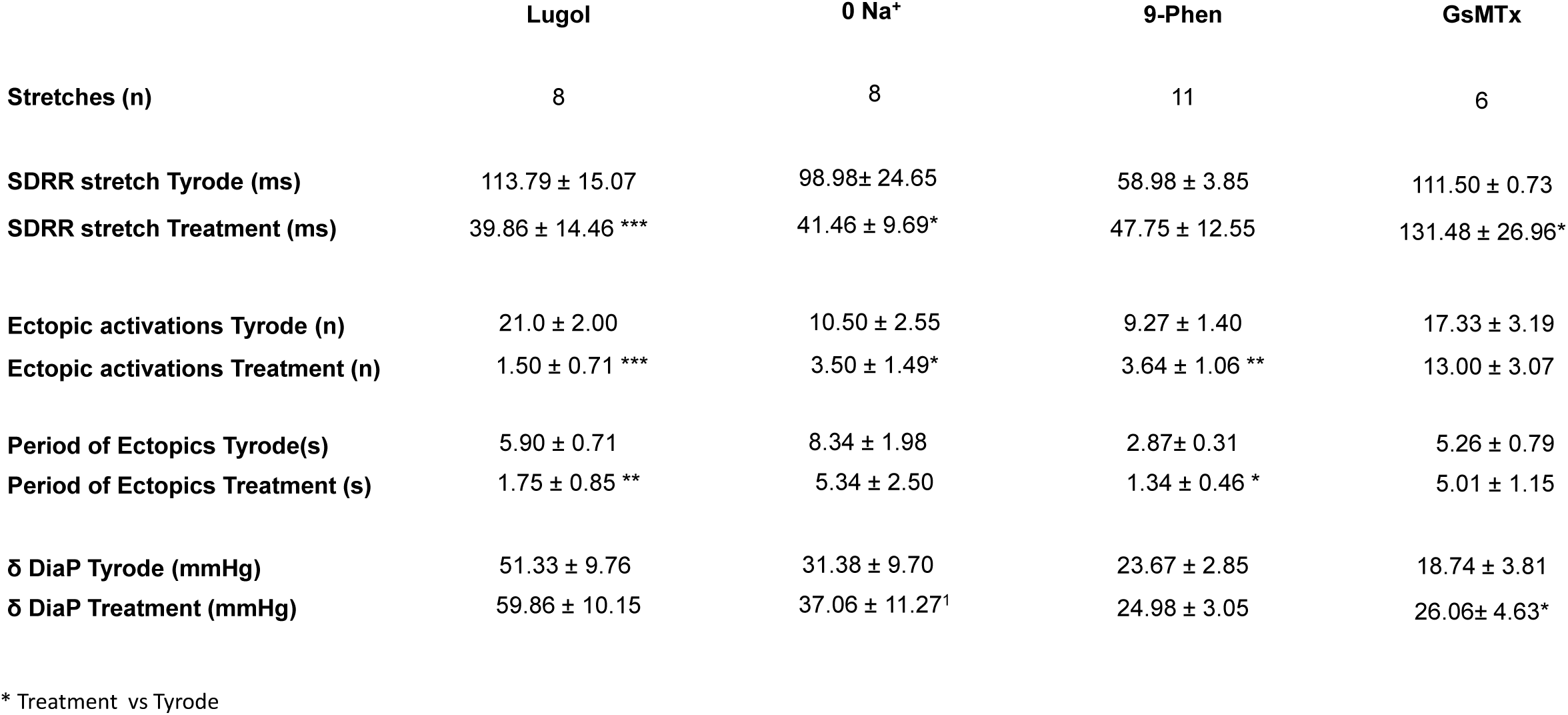
The effect of LV stretch following LV irrigation with Tyrode followed by irrigation with either Lugol’s (120mM KI, 38.6 mM I_2_), 0 Na^+^ Tyrode; 50μM 9-Phenanthrol or 1μM GsMTx4 upon: the standard deviation of the R-R interval, SDRR; the period of ectopics, the total number of ectopic activations and the change in diastolic pressure (an indicator of myocardial compliance, δDiaP). Lugol’s caused a statistically significant reduction in all 3 electrical parameters during stretch; 0 Na^+^ Tyrode significantly reduced the mechanically-induced increase in SDRR and the number of ectopic activations; 9-Phenanthrol significantly reduced the period of ectopics and number of ectopic activations but did not affect SDRR. GsMTx4 increased δDiaP and SDRR. Data are mean ± SEM, * P<0.05, ** P<0.01, *** P<0.001 paired t-test, Tyrode vs Treatment (SDRR, 2 way repeated measures ANOVA see Fig. 3).

**Fig 1.**
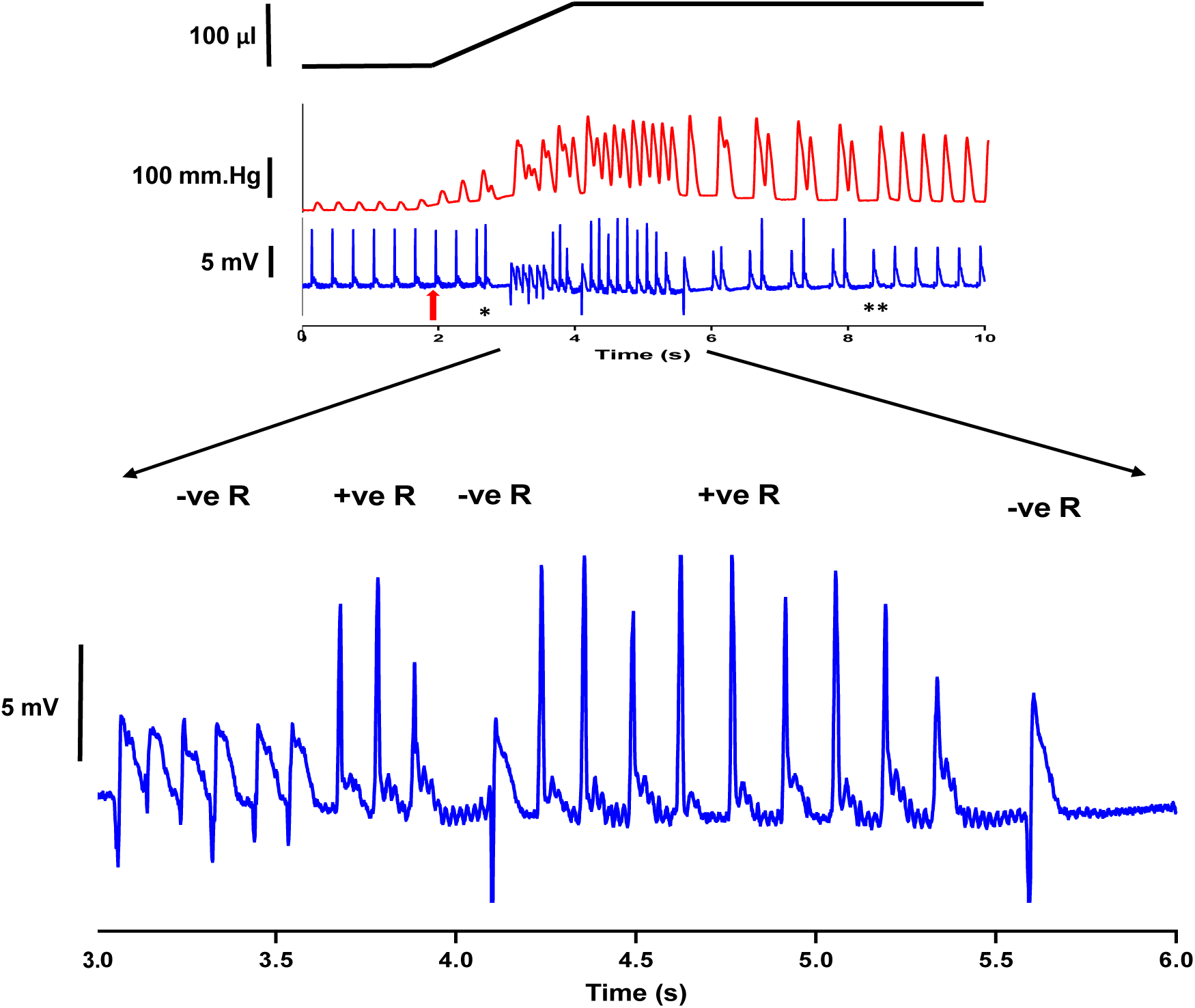
Upper trace, schematic of changes in balloon volume, changes in left ventricular (LV) pressure and pseudo-ECG from a Langendorff perfused isolated rat heart in sinus rhythm. **↑** (at approx. 2s) indicates the onset of LV stretch by injection of 100 μl into the balloon placed in the LV lumen. At approx. 2.5s * indicates the start and at approx. 8s ** indicates the end of mechanically-induced arrhythmias during the stretch and the subsequent spontaneous return to sinus rhythm. Sinus rhythm waveforms had stable but different waveforms prior to and during stretch. The change in waveform is likely caused by movement of the recording electrodes. Ectopic activations were identified as having a waveform different to either sinus configuration. The lower trace shows an expanded time scale of the pseudo-ECG trace highlighting a period of ectopic excitations with examples of –ve or +ve going initial waveforms.

Following brief endocardial exposure to Lugol’s solution, mechanically-induced arrhythmias were attenuated (abolished in the example given in Fig. 2A). When regular sinus rhythm was restored during stretch, the developed pressure was increased, consistent with an intact Frank-Starling mechanism (Fig. 2A). Visual inspection of hearts treated with Lugol’s solution confirmed the bulk of the LV myocardial wall retained a normal colouration at the end of the experiment (Fig 2B).

**Fig 2.**
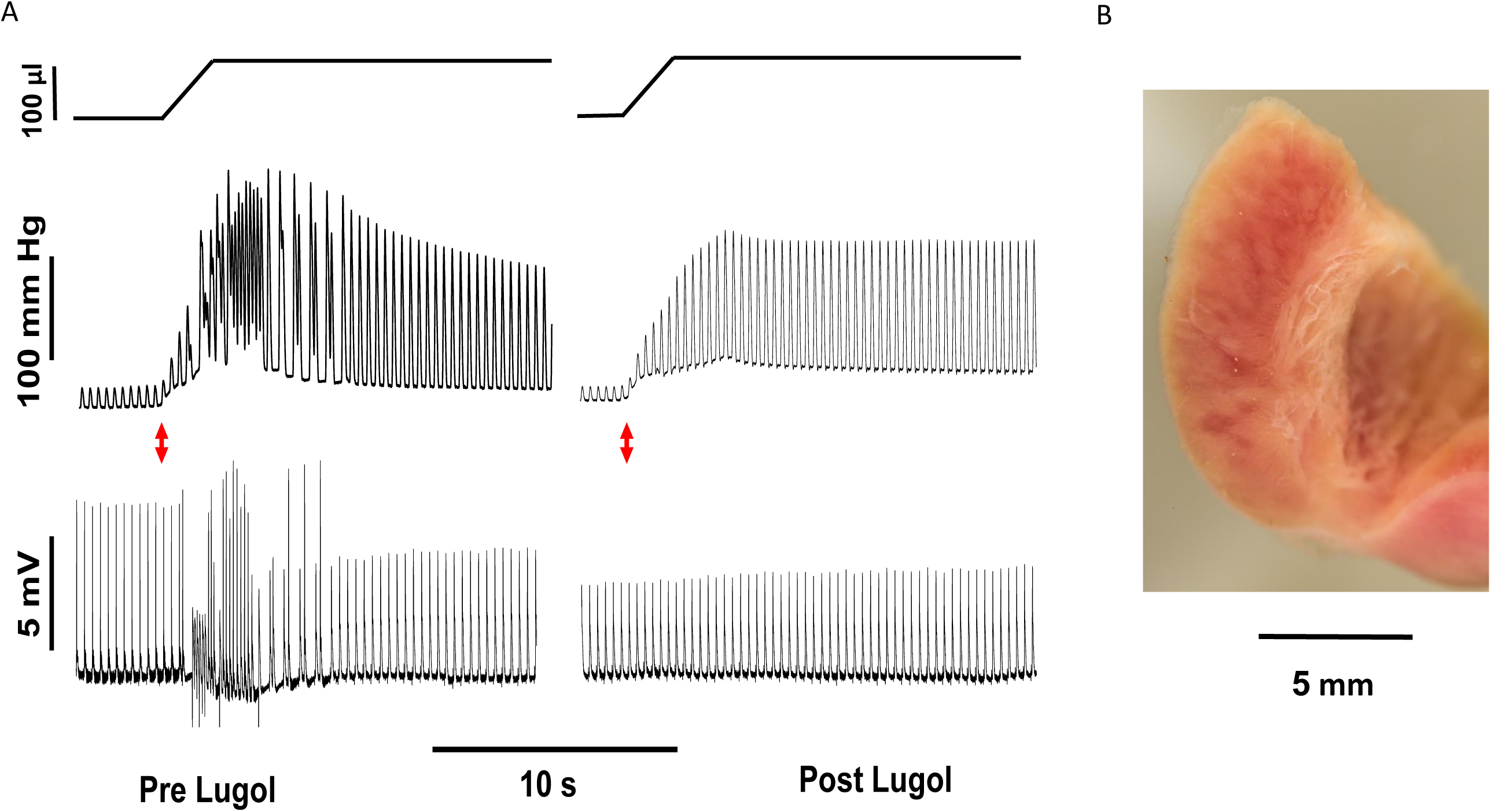
**A** Upper trace, schematic of changes in balloon volume, changes in left ventricular (LV) pressure and pseudo-ECG from a Langendorff perfused isolated rat heart in sinus rhythm. The red arrow indicates stretch of the LV by injection of 100μl into a balloon placed in the LV lumen. Records from the same heart before (Pre Lugol) and after (Post Lugol) irrigation of the LV lumen with 0.1ml Lugol’s over a 10s period. Mechanically-activated arrhythmias are evoked in the Pre Lugol traces, seen as disruption to the regular pressure transients and pseudo-ECG, R-waves. In this heart, Lugol’s prevented the development of mechanically-activated arrhythmias and Post Lugol the heart continues to beat rhythmically during stretch. The mechanical response of the heart remains intact, as indicated by the mechanically-induced increase in diastolic and systolic pressure (the Frank-Starling mechanism). **B** Section through a rat LV following an experiment involving Lugol’s treatment. Thin threadlike PFs are seen at the endocardial surface. This morphology explains their particular susceptibility to agents introduced into the LV lumen. The bulk of the LV myocardial wall retains a normal colouration.

Stretch significantly (P < 0.05) increased SDRR prior to all treatments (Fig. 3 Tyrode SDRR Pre vs Stretch) changes in SDRR within a given heart are shown in suppl. Fig S2. The mechanically-activated increase in SDRR was significantly reduced following Lugol and 0 Na^+^ (stretch SDRR: Tyrode vs Lugol’s, Fig 3A; Tyrode vs 0 Na^+^, Fig. 3B). Treatment with 9-Phen (Fig. 3C) did not attenuate the stretch-induced increase in SDRR (stretch SDRR Tyrode vs 9-Phen). Lugol’s, 0 Na^+^ and 9-Phen significantly reduced the number of ectopic activations and Lugol’s and 9-Phen significantly reduced the period of ectopic activity (see Table 1).

**Fig. 3.**
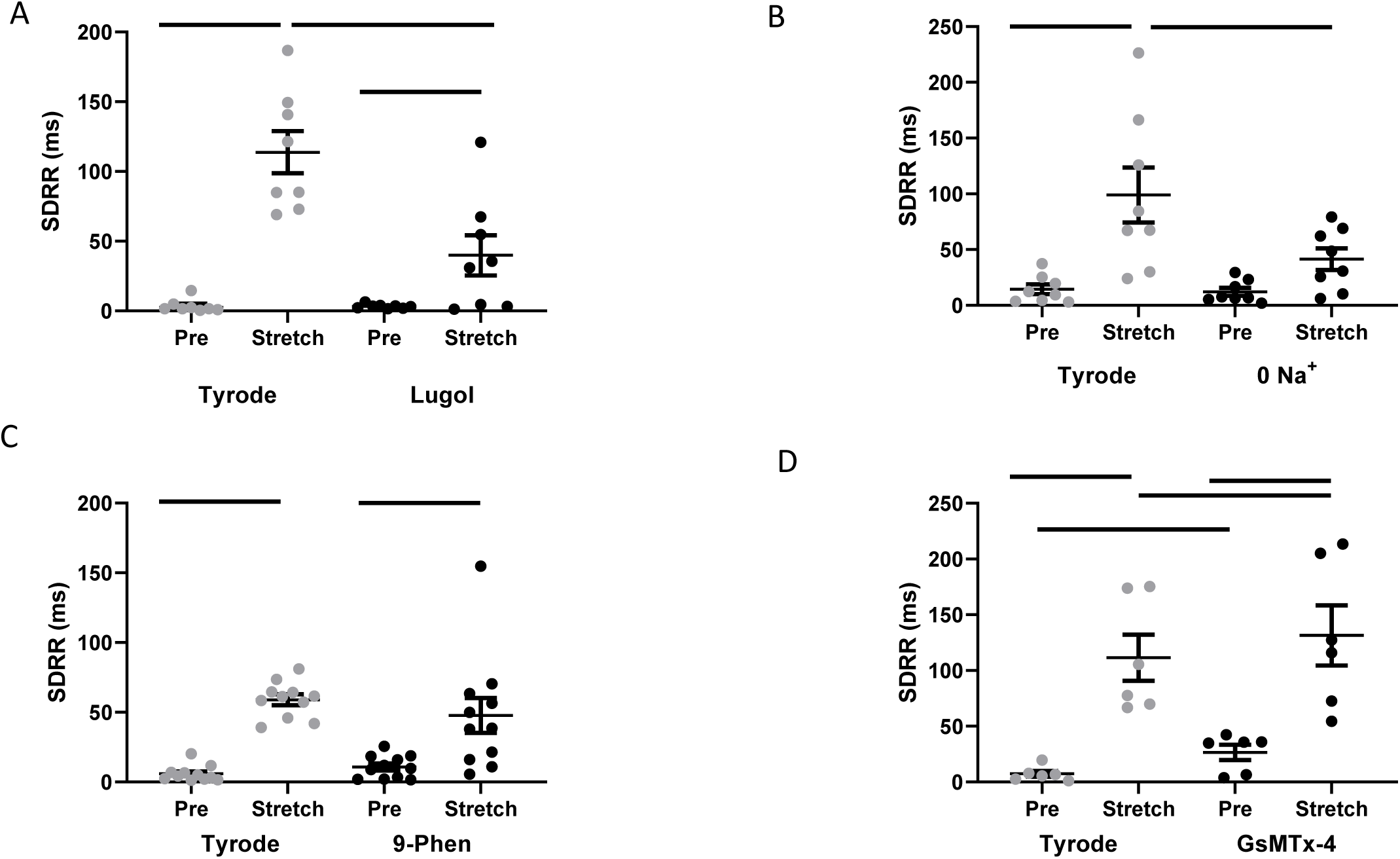
The standard deviation of R-R interval, SDRR, before (pre) and during (stretch) LV stretch following irrigation with Tyrode (Tyrode) and then either **A** Lugol’s (120mM KI, 38.6 mM I_2_), **B** 0 Na^+^ Tyrode; **C** 50μM 9-Phenanthrol or **D** 1μM GsMTx4. Stretch increased SDRR before each treatment. This increase was significantly attenuated by Lugol’s and 0 Na^+^. Data are individual datum and mean ± SEM. Bars indicate P<0.05 for post-hoc pairwise comparisons following 2-way repeated measures ANOVA, n = 6-11 stretches from N = 4-6 hearts in each group.

There was no effect of treatment on the mechanically-induced δDiaP in response to Lugol’s, 0 Na^+^ or 9-Phen, (Table 1), suggesting no effect upon myocardial compliance. Prior to stretch, there were no effects of treatment upon sinus heart rate. Only 9-Phen caused a significant change (increase) in pre-stretch LVDP (Table 2).

**Table 2.** Sinus heart rate (beats per min.) and left ventricular developed pressure (LVDP, mm.Hg) in isolated Langendorff perfused rat hearts, before and after LV irrigation with Lugol’s (120mM KI, 38.6mM I_2_), 0 Na^+^ Tyrode; 50μM 9-Phenanthrol or 1μM GsMTx4. The only significant effect was an increase in LVPD following 9-Phenanthrol. Data are mean ± SEM, * P<0.05, paired t-test Tyrode vs Treatment.

|  | Lugol | 0 Na<sup>+</sup> | 9-Phen | GsMTx |
| --- | --- | --- | --- | --- |
| Hearts (N) | 4 | 5 | 6 | 4 |
| Heart rate Tyrode (bpm) | 195 ± 15 | 193 ± 29 | 216 ± 22 | 178 ± 27 |
| Heart rate Treatment (bpm) | 220 ± 26 | 193 ± 16 | 203 ± 14 | 165 ± 22 |
| LVDP Tyrode (mmHg) | 12.59 ± 2.09 | 17.07 ± 8.53 | 11.07 ± 1.53 | 20.64 ± 8.80 |
| LVDP Treatment (mmHg) | 14.00 ± 2.57 | 20.96 ± 12.24 | 14.70 ± 2.66* | 25.22 ± 9.70 |

\* Treatment vs Tyrode

GsMTx4 did not affect intrinsic HR or LVDP (Table 2) but did increase δDiaP (Table 1) and appeared to decrease the stability of the preparations, causing a significant increase in SDRR both pre-stretch and during stretch (Fig. 3D). Stretch data from one heart treated with GsMTx4 was excluded as the heart became unstable prior to stretching. GsMTx4 had no significant effects on either the number or period of ectopic activations (Table 1).

In sheep LV preparations (Fig. 4A) focal mechanical stimulation of individual PFs provoked ectopic activations distinct from the activations triggered by electrical pacing of the endocardium (see Fig 4B). Optical mapping showed action potentials associated with both paced and ectopic beats (see upper trace Fig.4B). Ectopic activity was observed during periods of mechanical stimulation and did not align with the timing of the external electrical stimulation (see Fig 4B middle and lower traces) occurring in the diastolic periods between electrical activations. In isolated sheep LV preparations, spontaneous ectopic activations were rare. However, ectopic activations significantly increased during 10s periods of mechanical stimulation of PFs, in each of 4 LV preparations, when compared to the 10s period prior to mechanical stimulation (P<0.05). The mean data for the 4 preparations are given in Fig. 4C.

**Fig. 4.**
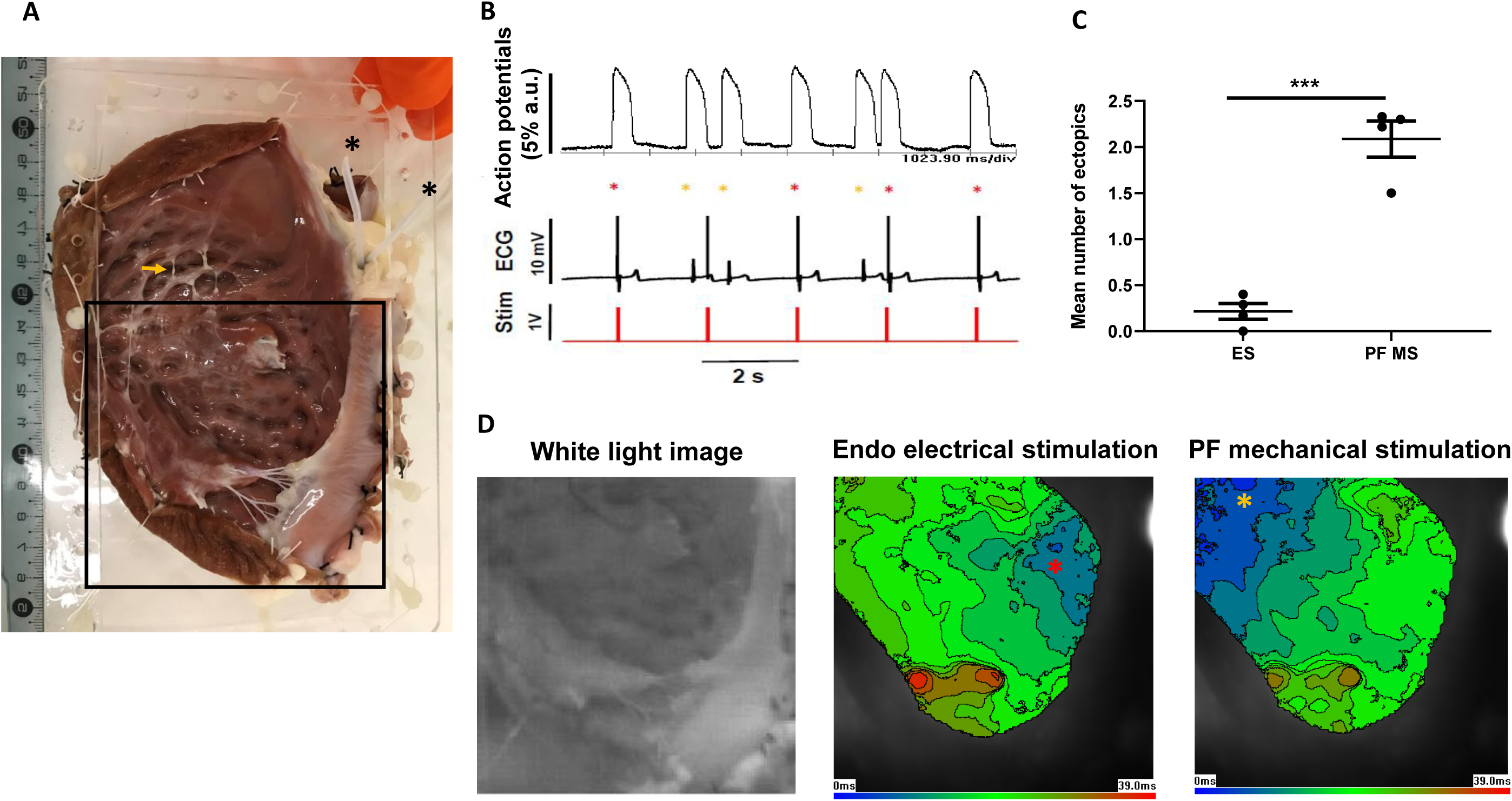
**A** Image of a sheep LV preparation prior to placement in the experimental chamber * indicate the 2 coronary cannulae. The orange arrow ↓ indicates the PF that was mechanically stimulated in this example. The square indicates the field of view for optical mapping in part **D**. Note that the 2 images do not superimpose due to the preparation taking up a more concave shape when immersed in the experimental chamber. **B** (upper trace) Optically mapped action potentials from the endocardial (ENDO) surface of the preparation shown in panel **A**. (Middle trace), pseudo-ECG and (lower trace) pacing electrode marker. * Indicates action potentials activated by electrical stimulation via pacing electrodes on the ENDO surface, note they align with the pacing marker. * Indicates action potentials activated during mechanical stimulation of the PF. The timing of the action potential and pseudo-ECG R-wave of ectopic activations do not align with the pacing marker and the form of the pseudo-ECG is distinct from paced activations. In this example there are 3 ectopic activations marked by * **C** Mean number of ectopic activations recorded in the 10 s interval before (ES) and during (PF MS) mechanical stimulation of single PFs (*** = P<0.001 paired t-test, N = 4 hearts). **D** Left, a white light image of the optically mapped region. Centre and Right, activation maps, showing the spatial and temporal distribution of the time of action potential upstroke, are shown for the first paced * and first ectopic * action potential in **A**. * and * are placed just below the position of earliest activation (blue 0 ms) which corresponds to the physical site of the pacing electrodes * or close to the mechanically stimulated PF *. The origin of these excitations are distinct and the electrical propagation (blue to green to brown/red) is in opposite directions. Isochrones at 3ms increments, field of view 8.5 × 8.5 cm, 850 * 850μm/pixel.

Mapping of electrical propagation in a sheep LV preparation is shown for electrical and mechanical stimulation (see Fig. 4D). The site of earliest activation (blue, 0ms) represents the origin of a given activation in the field of view. Green indicates intermediate time of activation (approx. 20ms) and brown/red the latest activation (approx. 40ms). A white light image of the optically mapped region is provided for orientation (see Fig. 4D). The central image represents the activation map associated with the first activation in Fig 4B, which is triggered by the pacing electrodes in the endocardium. As expected, the activation origin is at the site of the stimulating electrodes. The following action potential occurs during the diastolic period between electrical stimulations and begins close (approx. 1cm below) the site of the mechanically stimulated PF, spatially remote from the pacing electrodes. It can be seen that the propagation of activation from earliest to latest sites is in opposite directions in these 2 excitations (see Fig. 4D). PF-related activity was confirmed by electrical stimulation of the same PFs that were mechanically stimulated, which resulted in earliest activation in the same site as that for mechanical stimulation (Suppl Fig. S3).

## Discussion

### Main Findings

The main findings of our study are that in isolated whole rat hearts, mechanically-activated arrhythmias in the LV (generated by balloon inflation) were reduced by agents expected to preferentially target PFs, when these agents were superfused over the LV endocardial surface. We suggest that mechanically-induced arrhythmias provoked by balloon inflation can, at least in part, be triggered by mechanical stimulation of PFs. In support of these findings, mechanical stimulation of individual PF in isolated sheep LV increased the number of ectopic activations. Ventricular mechanically-activated arrhythmias have previously been thought to be a phenomenon of the myocardium. We now suggest that in addition to the myocardium, PFs may play a role in some mechanically-activated arrhythmias.

### Mechanically-induced arrhythmias in rat hearts

When considering the role of PF in the generation of mechanically-activated arrhythmias, it is of note that extensive endocardial surface PF networks have been identified in rat hearts e.g. (Bordas *et al*., 2010). They are particular susceptible to agents introduced into the LV lumen, because of their thin structure and surface location (Fig. 2B).

To quantitatively assess our interventions upon mechanically-induced arrhythmias we measured their number, their period and their impact on SDRR. SDRR is typically used as a measure of heart rate variability related to sinus node function over an extended period, but here it was calculated over 20s to give an amalgamated index of the number and complexity of ectopics. Pre-stretch SDRR was low (see Fig.3) but the generation of mechanically-induced arrhythmias transiently disrupted sinus rhythm and thus increased SDRR during-stretch.

We observed mechanically-activated ectopics of various types (see Fig. 1) and there was no consistent pattern to their format. Variation in the type of mechanically activated arrhythmia has been previously reported (e.g. Franz *et al*, 1992; Kohl *et al*, 2011; Quinn *et al*, 2017; Quinn & Kohl, 2020). In the rat LV, ectopics with a negative R-wave may originate from either the myocardium or the distal Purkinje network distant from origins affiliated with the normal sinus rhythm (e.g. the ventricular base). Positive waveforms may involve the proximal Purkinje network, in or near to the His bundle, enabling the global sinus rhythm pattern to be somewhat mimicked. Arrhythmias may have been sustained by re-entry or focal source mechanisms that involved the PF network, not necessarily to the exclusion of involvement of the myocardium.

### Mechanically-induced arrhythmias in sheep LV

Optical mapping cannot measure discrete activation within an *in situ*, single PF, rather the spread of activation into the coupled myocardium is measured. The activation map given in Fig. 4D in response to mechanical stimulation of a PF shows a broad activation pattern, consistent with rapid conduction through an extended portion of the distal Purkinje network (Martinez et al, 2018) and electrical stimulation of the same PF gave a similar activation pattern (Suppl. Fig. S3). Unknown variation may come from the number of parallel, but uncoupled PF bundles recruited within the stimulated PF branch by either electrical or mechanical modes. As we were unable to quantify the amplitude of the mechanical stimulus *in situ*, it was cautiously applied to avoid PF damage, this may explain why we only observed excitations in response to 20-25% of stimuli, compared to other studies that have been able to modify stimulus intensity and achieve 1:1 stimulus:activation e.g. (Franz et al, 1992; Quinn & Kohl, 2017).

### Local mechanical activation

Endocardial application of Lugol’s, 0 Na^+^ and 9-Phen, had no effect upon sinus heart rate, did not decrease LV developed pressure or change the mechanically-induced increase in diastolic pressure (an indicator of myocardial compliance). This suggests that their attenuation of mechanically-induced arrhythmias was via sources local to the endocardial surface, rather than the bulk of the myocardium. Local mechanical activation of PFs was demonstrated directly in sheep preparations. (Quinn *et al*., 2017) showed that in isolated rabbit hearts, focal mechanical stimulation of the epicardial surface could trigger arrhythmias arising from the epicardial site of mechanical stimulation. Both studies suggest that mechanically-induced arrhythmias arise from local sources rather than from the bulk of the myocardium.

### Site of action

Agents applied to the endocardial surface may affect the endocardial endothelium. Damage to the endocardial endothelium is known to affect the underlying myocardium (Brutsaert *et al*., 1996) and mechanical stimulation of the endocardial endothelium has been reported (Hoyer *et al*., 1994;Calaghan & White, 2001). However, the modulation of the myocardium by the endocardial endothelium is paracrine in nature and too slow (Brutsaert *et al*., 1996) to account for the rapid effects of stretch on electrical activity.

Another interpretation of our data is that arrhythmias were generated *exclusively* within the endocardial myocardium. The minimal effects of Lugol’s, 0 Na^+^ and 9-Phen on LVDP and stretch-induced δDiaP in rats suggest that if the stretch responses came exclusively from the endomyocardium, the area in question would need to be sufficiently deep to overcome sink-source issues, but thin enough that global mechanical properties were unaffected. To access the endocardial myocardium, agents must first cross, but not affect, the endothelium (see above). Furthermore, there is probably a steep fall in agent concentration at the endomyocardial surface due to diffusional limits into tissue that is also being perfused with Tyrode via the coronary circulation. This is relevant when interpreting the effects of 0 Na^+^ solution which was used to reduce general excitability and thus arrhythmic trigger and sustaining mechanisms, via reduction of phase 0 depolarisation.

If non-specific stretch-activated channel activation in the endocardial myocardium were responsible for the mechanically-induced arrhythmias, GsMTx4 would have been predicted to attenuate them. However, GsMTx4 did not reduce the number or period of ectopics and increased SDRR. One confounding factor is the increase in δDiaP in response to balloon inflation following GSMTx4 treatment. The data suggests a decrease in the stability of hearts following endocardial irrigation with GsMTx4 (increased pre-stretch SDRR and stretch-induced δDiaP). Instability is not reported when GsMTx4 is applied via the coronary circulation. Heterogeneities produced by selective irrigation of the LV lumen may explain these observations and may point to a role for non-specific stretch-activated channels in the normal function of the heart.

9-Phen, partially reduced, mechanically-activated arrhythmias. 9-Phen was thought to preferentially target PFs because of its action on TRPM4 channels. In species where the comparison has been made, TRPM4 is more abundant in conductive tissue than ventricular tissue (Liu et al., 2010;Kruse et al., 2009). TRPM4 single channel activity was reported in PFs but not in ventricular myocytes and 9-Phen modulated the action potential of PFs but not ventricular myocytes in rabbit (Hof et al, 2016). 9-Phen may target stretch-activation of TRPM4 channels, reported in arteries (Li et al., 2014; Earley et al., 2007) but as yet unstudied in cardiac PFs, or may cause a non-mechanically related generic decrease in excitability which increases the threshold for the triggering and/or sustaining of arrhythmias (Guinamard et al., 2015).

In sheep, direct mechanical stimulation of PFs *in situ* was demonstrated. When the PFs were contacted by the 2mm diameter probe there was little visible deformation of the underlying myocardium, where myocardium was present. Given the evidence (see Introduction) that PFs are mechanically sensitive it seems unlikely that PFs that were directly contacted were mechanically inert but that myocardium was mechanically activated by the transfer of a damped stimulus via the PF.

### Mechanism of action

Lugol’s is a well characterised and accepted PF ablative agent and we found this agent most effective at attenuating mechanically-induced arrhythmias. However it did not abolish arrhythmias in all preparations. Therefore arrhythmias arising in or sustained by the myocardium may form part of the response we saw. We do not expect the roles of the PF network and the myocardium, in mechanically-induced arrhythmias, to be mutually exclusive.

Ventricular dilation may cause an increase in PF strain and stress. Previous work (Dominguez & Fozzard, 1979, Sanders et al, 1979 Canale et al, 1983) has suggested that PFs can extend by a combination of fibre unbuckling and sarcolemmal unfolding, such that the increase in membrane strain is less than the increase in fibre length. These properties may explain the high compliance of PFs relative to the myocardium and the observation that studies reporting modulation of electrical activity in isolated PFs do so typically at greater extensions (e.g. Saunders et al 1997, 40%; Kaufman & Theophile 1967, 40-70%; Dominguez & Fozzard et al 1979, 30-50%) than those typically reported for the myocardium (around 20% see studies in Kohl et al, 2011). Tissue deformation caused by the physical contact between the balloon and rat endocardial surfaces or by the probe and sheep PF surfaces may also be important. Electrical activation upon surface deformation with a probe has been demonstrated and characterised in isolated myocytes e.g. Betts & Sachs, (2000) and in intact hearts e.g. Quinn et al, (2017).

The (almost) 2D cable properties of thin PFs reduce sink-source issues associated with electrical activation and conduction, when compared with the myocardium (Haissaguerre *et al*., 2016) this may be an important factor in the response of the Purkinje network to mechanical stimulation. In addition to the PFs, the Purkinje system includes Purkinje-Muscle junctions. These junctions contain specialised transitional cells that are a source of conduction delay and repolarisation heterogeneities (Wiedmann et al, 1996). The mechanical stimulus applied to the PF may be transferred to the Purkinje-Muscle junction. Furthermore, retrograde conduction from the myocardium into the Purkinje system is a factor in some arrhythmias (Haissaguerre et al, 2016). Any or all of these pathways may be involved in the role of the Purkinje system in mechanically-activated arrhythmias and involve both triggering and sustaining mechanisms. The aim of this study was to provide evidence for the involvement of the Purkinje system in mechanically-activated arrhythmias, further studies are required to investigate the above possibilities.

### Study limitations

We did not inspect the effect of Lugol’s by histology, however Lugol’s, is a well established treatment for chemical PF ablation e.g. (Chen *et al*., 1993; Dosdall *et al*., 2008;Myles *et al*., 2012) and we observed no effects of Lugol’s on heart rate or pressure development (Table 2). 9-Phen was reported, to target late Na^+^ current in some pathologies (Hou *et al*., 2018). However, late Na^+^ current is not a large component of normal rat electrophysiology e.g. (Brette & Orchard, 2006). It can become important if increased under pathological conditions (Saint, 2006). The study by (Hou et al 2018) shows little effect of 9-Phen in normal tissue but large effects when the late Na^+^ current is enhanced by ATXII or by overexpression in CHO cells. Conditions that enhance late I_Na_^+^ did not apply to our experiments. We have not directly measured the presence or altered activity of TRPM4 in rat PFs. The presence of TRPM4 (mRNA, protein and current) has been confirmed in rat ventricular tissue (Guinamard *et al*., 2006;Guinamard, 2007;Piao *et al*., 2015). There is direct evidence for the presence of TRPM4 in rabbit (Hof *et al*., 2016) and bovine (Liu *et al*., 2010) PFs and indirect evidence for its presence in the conductive tissue of rat (Hof *et al*., 2013). In species where the comparison has been made, TRPM4 is more abundant in conductive tissue than ventricular tissue (Liu *et al*., 2010;Kruse *et al*., 2009) see (Guinamard *et al*., 2015) for review. Currently, quantification of the timing of the mechanical stimulus applied to sheep PFs is lacking.

### Potential implications

PFs are recognised as an important factor in clinical arrhythmias e.g. (Haissaguerre *et al*., 2016) as are mechanically-induced arrhythmias (Orini *et al*., 2017; Quinn & Kohl, 2020). These 2 factors may interact in pathological settings where mechanical and electrical heterogeneities within and between PFs, Purkinje-Muscle junctions and myocardial tissue may be increased, for example, at the border zone of an infarct (Ferrier, 1976) in ischemic tissue (Gilmour et al, 1984), in mitral valve prolapse (Fulton et al 2020) or dilated cardiomyopathy (Sinha et al, 2009).

## Acknowledgements

Supported by The Universities of Auckland, Leeds and Bordeaux, The European Union (Marie Curie International Research Staff Exchange Scheme CORDIS-3D) and British Heart Foundation grant PG/19/47/34335. The authors thank Mr. Dane Gerneke for preparation of Lugol’s solution, Ms. Sasiththika Arulanantham, Ms. Kayleigh Parslow and Ms. Anna Maria Krstic for experimental assistance and Ms. Virginie Loyer for expert animal technical services.

## Supplementary Materials

**Figure S1.**
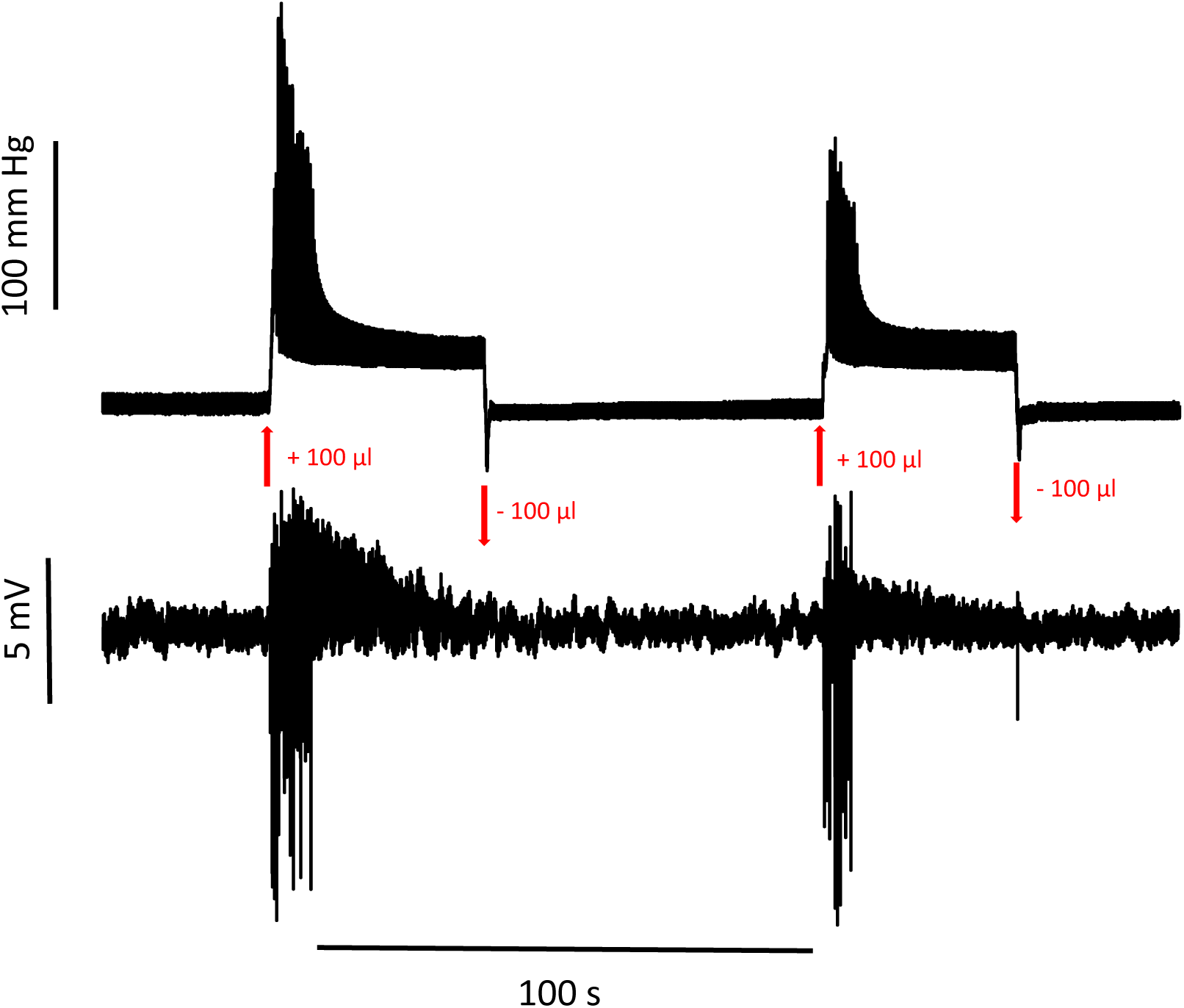
Continuous recording of changes in left ventricular (LV) pressure (upper trace) and pseudo-ECG (lower trace) from a Langendorff perfused isolated rat heart in sinus rhythm. Arrows indicate the inflation and deflation of a balloon placed in the LV lumen. The inflation/deflation cycle is repeated twice. At this timescale individual pressure transients and activations cannot be identified.

**Figure S2.**
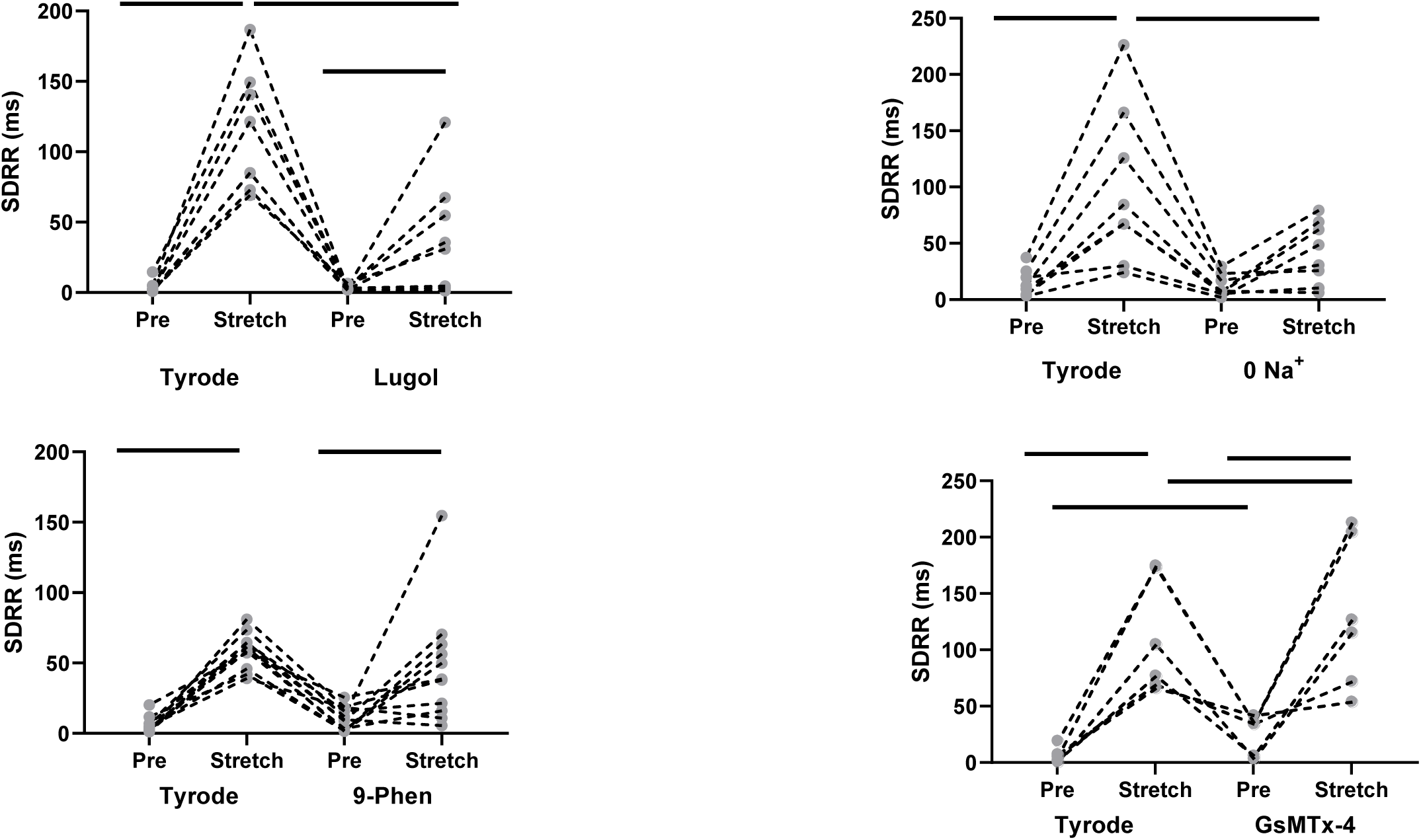
The standard deviation of R-R interval, SDRR, before (pre) and during (stretch) LV stretch following irrigation with Tyrode (Tyrode) and then either **A** Lugol’s (120mM KI, 38.6mM I_2_), **B** 0 Na^+^ Tyrode; **C** 50μM 9-Phenanthrol or **D** 1μM GsMTx4. Stretch increased SDRR before each treatment. This increase was significantly attenuated by Lugol’s and 0 Na^+^. Data are individual datum and dotted lines connect observations from a given heart, Bars indicate P<0.05 for post-hoc pairwise comparisons following 2-way repeated measures ANOVA, n = 6-11 stretches from N = 4-6 hearts in each group. (see Fig. 3).

**Figure S3.**
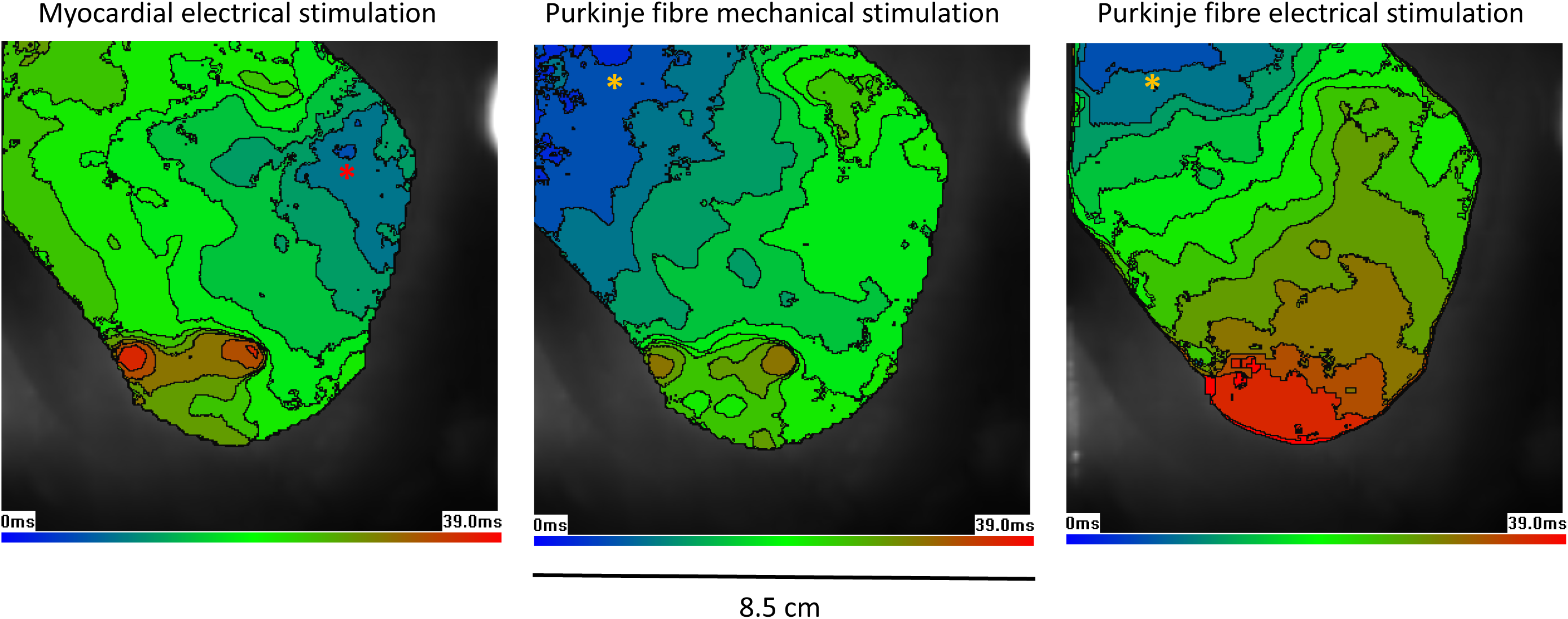
Activation maps, showing the spatial and temporal distribution of the time of action potential upstroke. **A** represents activation by electrical stimulation of the myocardium **B** represents the subsequent ectopic activation during mechanical stimulation of a PF. **C** represents activation in response to electrical stimulation of the same PF. * are placed just below sites of earliest activation (blue). In **A** this corresponds to the physical location of the stimulating electrodes in **B** & **C** this is approx. 1cm below the PF. The origin of the excitation and direction of conduction (blue to green to brown/red) in **B** & **C** are the same and distinct from **A**. Isochrones at 3 ms increments, field of view 8.5 × 8.5cm, 850 * 850μm/pixel.

